# Root Water Uptake and Photosynthesis Synergistically Enhance Drought Tolerance in Interspecific Hybrid Sugarcane

**DOI:** 10.1101/2025.07.01.662668

**Authors:** Naoya Katsuhama, Shotaro Tamaru, Shin Yabuta, Wataru Yamori, Jun-Ichi Sakagami

## Abstract

Sugarcane (Saccharum spp. hybrids) is a globally important crop for food and bioenergy, but its production is increasingly threatened by drought driven by climate change. Drought causes complex interactions between root function and leaf photosynthesis, but their underlying mechanisms remain largely unstudied in sugarcane. To address this knowledge gap, we grew up to eight sugarcane cultivars in one field and two pot experiments inside a rain-out shelter. The relative growth rate of these eight cultivars under both well-watered and water-limited conditions was largely explained by net assimilation rate, which depended on bleeding sap rate caused by root pressure. Among the cultivars, Harunoogi—a first-generation backcross cultivar between a wild relative (S. spontaneum) and a commercial cultivar—exhibited improved drought avoidance through adequate root water supply, whereas the commercial cultivar NiTn18 showed the opposite. Throughout the drought and recovery, Harunoogi exhibited significantly higher leaf area, stomatal conductance, chlorophyll content, and maximum amplitude of the reaction center chlorophyll of PSI than cultivar NiTn18, which resulted in a higher net photosynthetic rate and biomass. These results highlight the potential of improving both root function and photosynthesis as an effective approach to developing drought-tolerant sugarcane.

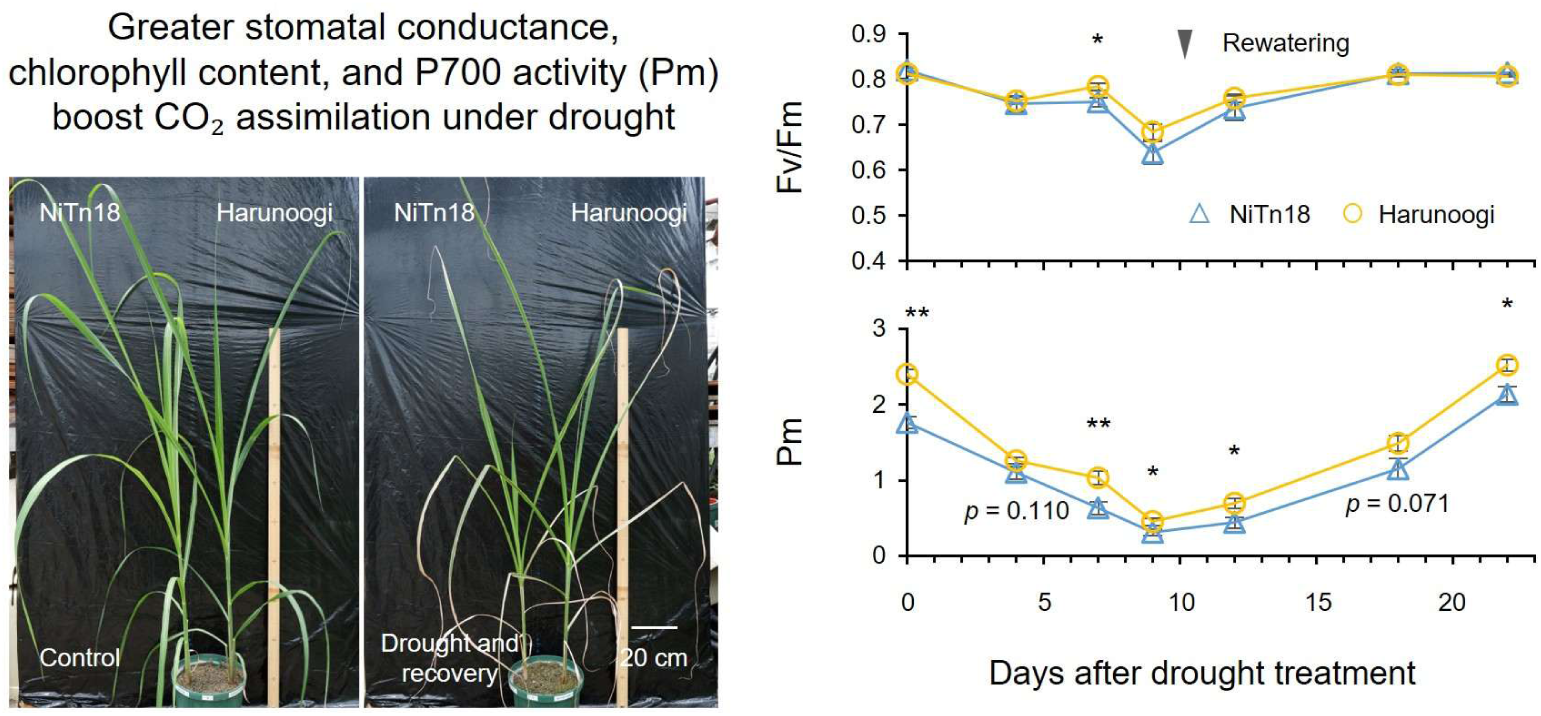

## Introduction

Sugarcane (*Saccharum* spp. hybrids)—the most harvested crop worldwide by tonnage (FAO, 2024)—is an economically important C_4_ plant grown globally for food and bioenergy. In 2023, sugarcane accounted for 87% of the global sugar crop production and 24% of bioethanol output, with a projected growth of 1% annually through 2033 (OECD/FAO, 2024). However, drought remains a critical threat to sugarcane yields, as reported in major sugarcane-producing countries such as Brazil (Cursi et al., 2022), India (Mall et al., 2022), and Australia (N. G. Inman-Bamber et al., 2012). Considering that drought frequency and severity are expected to increase due to climate change (IPCC, 2022) and that agricultural water availability is projected to decline by 2050 (UNESCO, 2020), developing drought-tolerant cultivars is a global priority.

Crop yield potential depends on three components: canopy light interception, conversion of intercepted light into biomass (i.e., photosynthesis), and harvest index (G. Inman-Bamber et al., 2013). Under agricultural drought, reduced water uptake from roots limits nutrient supply, leaf expansion, and photosynthesis, leading to yield losses (Vadez et al., 2024). In sugarcane, culm elongation and canopy development are especially sensitive to water deficits, followed by photosynthesis (Lakshmanan & Robinson, 2013). Conversely, sucrose yield remains closely tied to aboveground biomass, even when growth declines by up to 50%, suggesting that the harvest index is stable under water stress (Robertson et al., 1999). This is not the case in grain crops, where reproductive development is highly sensitive to drought (Vadez et al., 2024). Thus, enhancing whole-plant photosynthesis is considered a promising strategy for improving sugarcane yield stability under drought conditions (G. Inman-Bamber et al., 2013; Basnayake et al., 2015). Specific root traits support carbon assimilation under water-limited conditions by enabling water and nutrient uptake (Vadez et al., 2024; Katsuhama et al., 2025). Previous studies have identified significant phenotypic variation in root traits among crops such as rice (Uga et al., 2013), maize (X. Wang et al., 2016), and grain legumes (Ye et al., 2018) exposed to drought conditions. For example, in rice, deep-rooting associated with a steeper root growth angle has been shown to improve drought avoidance, maintaining stomatal conductance (*g*_sw_), net CO_2_ assimilation rate (*A*), and yield (Uga et al., 2013). Similarly, maize with a higher number of roots and larger root biomass exhibited higher *g*_sw_, *A*, and yield under drought stress (X. Wang et al., 2016). In sugarcane and related species (*Saccharum* complex), there can be significant natural variation in root growth under drought (Augustine et al., 2015; Chanaphai et al., 2023; Terajima et al., 2023).

Considering that the genetic base of commercial sugarcane is narrow due to its limited progenitors, interspecific and intergeneric hybridization within the *Saccharum* complex is gaining attention as a strategy to improve both productivity and stress resilience (Hattori et al., 2019; Terajima et al., 2022). Notably, an intergeneric hybrid between sugarcane and *Erianthus arundinaceus*, a related germplasm characterized by deep roots and high drought tolerance, was shown to develope deeper roots than its parental sugarcane cultivar (Terajima et al., 2023). Furthermore, *g*_sw_ and *A* vary widely within the *Saccharum* complex under drought conditions (Augustine et al., 2015; Basnayake et al., 2015; P. Jackson et al., 2016; Endres et al., 2019; Takaragawa & Wakayama, 2024). Moreover, a simulation study identified increased root vigor and leaf photosynthesis as key traits for improving sugarcane yield in water-limited environments (N. G. Inman-Bamber et al., 2012). However, comprehensive studies linking root function and photosynthesis remain limited.

Photosystem II (PSII) and photosystem I (PSI) are key components of the photosynthetic electron transport system, which converts light energy into reducing power for carbon fixation. Photoinhibition of PSI and PSII depends on the type, duration, and severity of abiotic stress (Yamori, 2016; Hu et al., 2023). Most investigations have focused on the photoinhibition of PSII, with field studies showing variation in it within the *Saccharum* complex (de Almeida Silva et al., 2011; Endres et al., 2019). In contrast, PSI inhibition is less studied, but it has been shown that this system is susceptible to fluctuating light in natural environments (for reviews, see Yamori and Shikanai, 2016; Kono et al., 2024). Moreover, recent findings indicate that PSI is particularly vulnerable not only under fluctuating light but also under drought conditions, both of which lead to constraints on carbon fixation (Sun et al., 2023; X.-Q. Wang et al., 2024). Despite these findings, no study has systematically analyzed the interactions between the photosynthetic electron transport system (PSI and PSII) and leaf gas exchange in sugarcane under drought stress.

In this study, we evaluated natural variation in shoot growth and root water uptake across eight sugarcane cultivars exposed to drought conditions by combining a simple measurement of sap flow in the root xylem with classical growth analysis. Additionally, we explored the correlation between these variations and leaf-level gas exchange to elucidate the relationship between root traits and photosynthesis. Furthermore, we investigated how drought responses in PSI and PSII influence overall photosynthetic performance. The aim of these analyses was to develop a novel strategy for improving sugarcane’s drought tolerance by simultaneously enhancing both root and leaf functions.

## Materials and Methods

### Study Site and Plant Materials

Experiment 1 was conducted in a sloped field (Figure S1) inside a rain-out shelter at Kagoshima University, Japan (31°34’23“;N, 130°32’34”E), whereas Experiment 2 was pot experiment. Experiment 3 replicating Experiment 2 was carried out at the University of Tokyo, Japan (35°44’13“N, 139°32’21”E) using the same design. Environmental data [i.e., solar irradiance, air temperature, relative humidity, and soil volumetric water content (VWC)] were recorded hourly. Soil VWC was measured using a 5TE sensor (METER Group, USA) at depths of 10 cm and 30 cm in Experiment 1 and at 10 cm in Experiment 2.

The following eight sugarcane (*Saccharum* spp. hybrid) cultivars common in Japan (Terajima et al., 2022) were selected for the experiments: NCo310, NiF8, NiTn18, Ni22, Ni23, Ni27, KN00-114, and Harunoogi. Since most global sugarcane cultivars share limited progenitors (P. A. Jackson, 2005; Terajima et al., 2022), our findings may be widely applicable to sugarcane cultivars worldwide. Single-bud setts with a length of 5 cm were soaked in a Benlate solution (5 g L^−1^; Sumitomo Chemical, Japan) for 12 h to ensure sterilization and promote germination. Seedlings were grown in a cell tray (0.15 L per cell) for 40–50 days until reaching the leaf age of 3–4. All tillers were removed immediately after emergence to eliminate the effects of tillering among cultivars.

### Drought Treatment

Experiment 1 was conducted in a terraced concrete container filled with Andosol and sand at a ratio of 1:2 (v/v) and divided into four plots with different soil surface elevations, i.e., 35 cm, 55 cm, 75 cm, and 95 cm above the groundwater level, to create a soil moisture gradient (Figure S1). The eight selected cultivars were randomly transplanted at a 15 × 15 cm spacing, and the planting was repeated in two fields as replicates. Chemical fertilizer (14.4 g m^−2^ N, P_2_O_5_, and K_2_O) was applied uniformly to the surface soil (top 10 cm). All plots were irrigated from above for 10 days, and treatments were applied for 14 days by switching to bottom watering only.

Based on the results of Experiment 1, cultivars NiTn18 (drought-sensitive) and Harunoogi (drought-tolerant) were used for Experiments 2 and 3. These two cultivars were transplanted into the same 2.25-L pots with 10-cm spacing to ensure equivalent soil water conditions. The soil, consisting of Andosol mixed with sand at a ratio of 1:2 (v/v), was added with chemical fertilizer (0.24 g pot^−1^ N, P_2_O_5_, and K_2_O). Drought was imposed by withholding water for 12 days (Experiment 2) or 11 days (Experiment 3), followed by rewatering.

### Growth Analysis

Leaves were collected and their area was measured (AAM-9, Hayashi Denko, Japan). Then, harvested leaves and stems were oven-dried at 80°C for 72 h and weighed. The mean values of aboveground growth analysis parameters during drought treatment were calculated using the classical approach (Hunt et al., 2002).

### Bleeding Sap Rate

In Experiment 1, root xylem sap flow caused by root pressure was measured shortly after dawn as described (Sakaigaichi et al., 2007). In brief, after cutting the stem at 10 cm above the ground, the section was covered with pre-weighed cotton and sealed with flexible plastic film (Parafilm M, Amcor, USA). After 1 h, the cotton was reweighed to calculate the bleeding sap rate. The leaf number was determined as an indicator of phytomer count to then estimate the root number (Kato et al., 2007). Subsequently, the bleeding sap rate per plant was divided by the leaf number to calculate water uptake per root (Morita & Abe, 2002).

### Leaf Gas Exchange and Chlorophyll Fluorescence of PSI and PSII

In Experiment 2, steady-state gas exchange was measured using an LI-6400 equipped with a 6400-02B chamber (LI-COR, USA). All measurements were conducted on the central portion of the youngest fully expanded leaves between 07:00 and 13:00. The chamber conditions were set at a photosynthetically active photon flux density (PPFD) of 2000 µmol m^−2^ s^−1^, air temperature of 30°C, relative humidity of 65% (air VPD of 1.48 kPa), and ambient CO_2_ concentration ([CO_2_]) of 400 µmol mol^−1^. The maximum quantum yield of PSII (Fv/Fm) was measured after overnight dark adaptation using AquaPen AP 100-P (PSI, Czech Republic).

In Experiment 3, before drought treatment, gas exchange and chlorophyll fluorescence were measured using an LI-6800 equipped with a 6800-01A chamber (LI-COR, USA) at a PPFD of 2000 µmol m^−2^ s^−1^. Measurements were taken at reference [CO_2_] of 400, 50, 100, 200, 300, 400, 600, 800, 1000, 1200, and 1500 µmol mol^−1^.

Rectangular saturation pulses (10000 µmol m^−2^ s^−1^, 1000 ms) were applied after overnight dark adaptation and at each [CO_2_]. The quantum yield of PSII (Y(II)), non-photochemical quenching (NPQ), and the fraction of open PSII reaction centers according to the Lake model (qL) were calculated (Kramer et al., 2004). The apparent maximum rate of carboxylation by PEPC (*V*_pmax_) was estimated using *A*/*C*_i_ curves (von Caemmerer, 2021). During drought and recovery, the maximum quantum yield of PSII (Fv/Fm) and the maximum amplitude of P700 (Pm) were simultaneously measured after overnight dark adaptation using the DUAL-PAM-100 (Heinz Walz, Germany) (Klughammer & Schreiber, 2008). A rectangular saturation pulse of 24412 µmol m^−2^ s^−1^ (P. + F. SP intensity of 20) was applied for 800 ms.

### Chlorophyll and Carotenoid Contents

Chlorophyll content in the same leaves used for gas-exchange measurements was non-destructively estimated using a SPAD-502 (Konica Minolta, Tokyo, Japan). Leaf disks (3 cm^2^) were subjected to extraction using 3 mL of *N,N*-dimethylformamide at 4°C in the dark for 72 h. Absorbance was determined using a spectrophotometer (GENESYS 50, Thermo Scientific, USA) at 480.0, 646.8, and 663.8 nm. Chlorophyll *a* and *b* and total carotenoid (Car *x* + *c*) contents per leaf area were determined as described (Wellburn, 1994).

### PEPC, PPDK, and Rubisco Contents

Leaf disks (3 cm^2^) collected from the same leaves used for gas exchange measurements were immediately frozen in liquid nitrogen. Total proteins were extracted and separated by SDS-PAGE as described (Yamori et al., 2011). Protein bands corresponding to phospho*enol*pyruvate carboxylase (PEPC), pyruvate, phosphate dikinase (PPDK), and the large subunit of Rubisco (RbcL) were stained with colloidal Coomassie dye G-250 (24590; Thermo Fisher Scientific, USA) and quantified using the Gel Analyzer tool in ImageJ/Fiji software (Schindelin et al., 2012).

### Leaf Mass per Area and Total Carbon and Nitrogen Contents

Leaf disks (3 cm^2^) collected from the same leaves used for gas exchange measurements were dried, weighed, and ground. Total C and N contents (%) were measured using an NC analyzer (SUMIGRAPH NC-22F; Sumika Chemical Analysis Service, Japan).

### Stomatal Traits

Stomatal complex density and length (over 20 stomata examined per biological replicate) were measured using an electron microscope (JCM-6000, JEOL, Japan). All measurements were conducted on both the adaxial and abaxial surfaces of the central portion of the youngest fully expanded leaves.

### Statistical Analysis

Sample sizes and statistical tests are included in the figure legends. Data were analyzed by Student’s t-test or two-way ANOVA with Tukey’s test at *p* < 0.05 using R v. 4.4.0 software (https://www.r-project.org/).

## Results

### Experiment 1: Shoot Growth under Well-Watered and Drought Conditions was Driven by Net Assimilation and Root Water Uptake

We examined the relationship between root function and shoot growth in the eight sugarcane cultivars subjected to varying water availability in the field experiment (Figure S1). Over the 14-day treatment, soil VWC at a depth of 10 cm in plot 1 ranged from 0.16 to 0.18 m^3^ m^−3^, whereas that in plots 2, 3, and 4 it gradually declined to 0.12, 0.09, and 0.06 m^3^ m^−3^, respectively (Figure S2a). As soil VWC decreased, shoot dry weight, leaf area, leaf number, and bleeding sap rate declined as well, with marked differences among cultivars (Figure 1). Notably, in plot 1, both NiTn18 and Harunoogi exhibited the highest shoot growth and root sap flow (Figure 1a, d). In contrast, in plot 4, Harunoogi maintained 80.8% of its shoot dry weight and 42.0% of its bleeding sap rate, whereas the proportions retained by NiTn18 were only 59.7% and 16.2%, respectively (Figure 1a, d).

**Figure 1.**
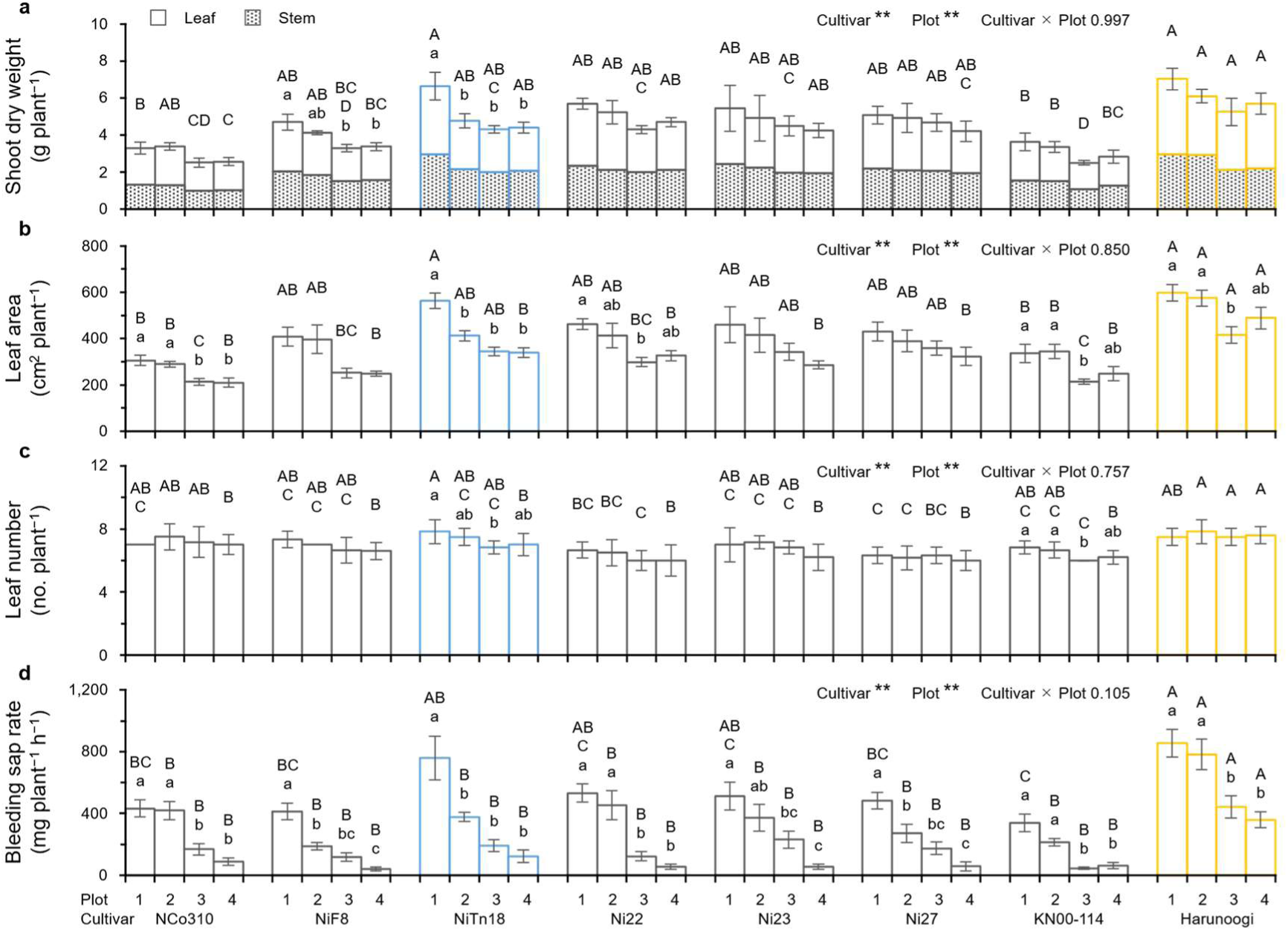
Shoot growth and root water uptake in eight sugarcane cultivars under four water conditions. **a** Shoot dry weight; **b** leaf area; **c** leaf number; and **d** bleeding sap rate per plant. Error bars represent SEM (*n* = 5–6). Mean values not sharing any letters are significantly different between cultivars (uppercase) or plots (lowercase) at *p* < 0.05 by Tukey–Kramer test. Significant main effects of cultivars or plots are denoted as \*\**p* < 0.01 by two-way ANOVA.

With regard to the relationship between growth analysis parameters and bleeding sap rate, we found a linear relationship between the mean relative growth rate (RGR) during the treatment and the mean net assimilation rate (NAR) explaining 84.7% of the variation (Figure 2a). This indicates that changes in shoot growth largely reflect the net photosynthetic carbon gained through photosynthesis minus the carbon lost through respiration. Furthermore, bleeding sap rate explained 58.8% of the variation in NAR across drought treatments (Figure 2d).

**Figure 2.**
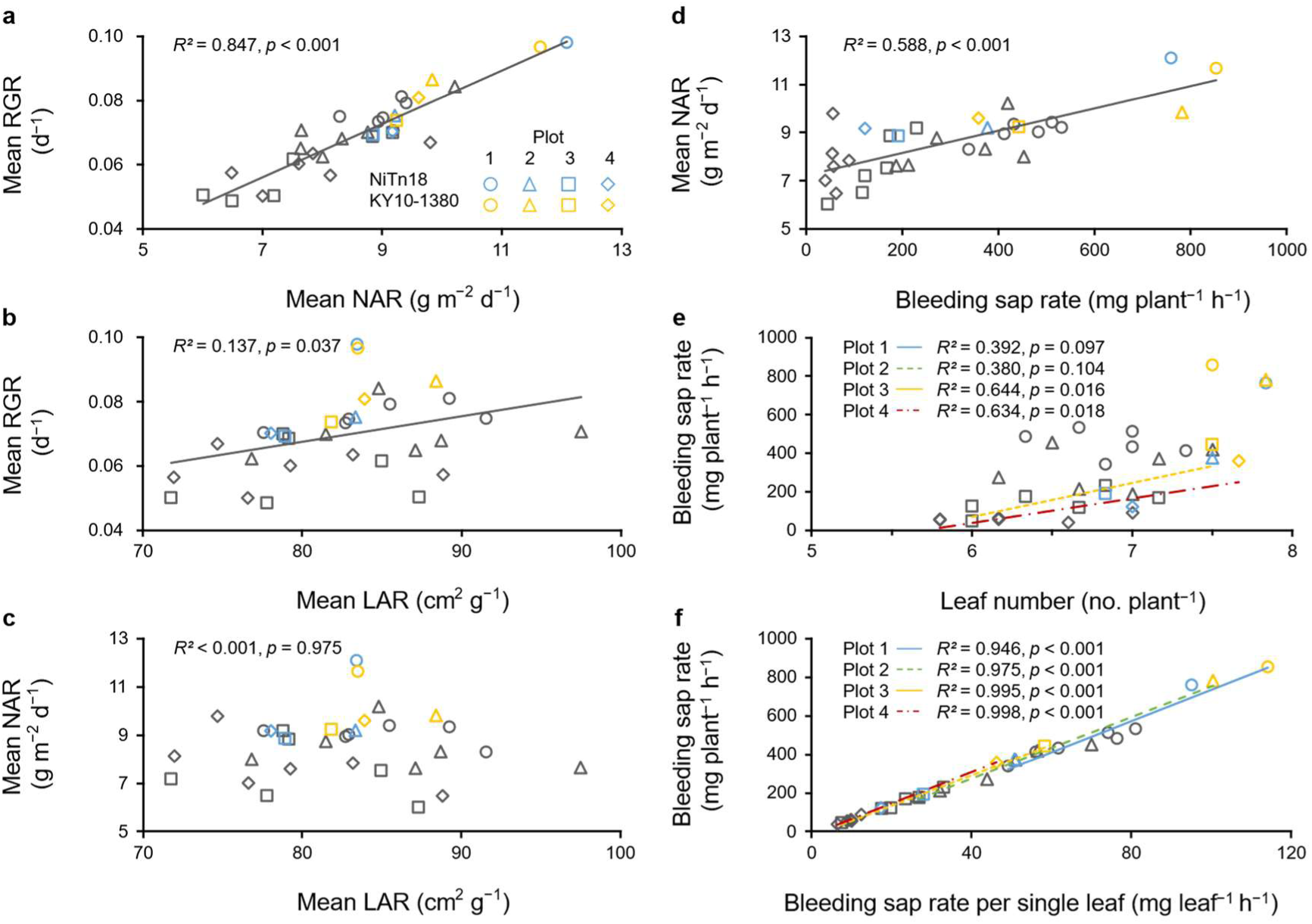
Relationship between mean relative growth rate (RGR), mean net assimilation rate (NAR), mean leaf area ratio (LAR), and bleeding sap rate under four water conditions. **a** Mean RGR and NAR; **b** mean RGR and LAR; **c** mean NAR and LAR; and **d** mean NAR and bleeding sap rate per plant. Indices were calculated using mean shoot dry mass and leaf area for 5–8 biological replicates before and after treatment (*n* = 32). Bleeding sap rate was measured at the end of the treatment in 5–6 biological replicates. **e** Bleeding sap rate per plant and leaf number; **f** bleeding sap rate per plant and per single leaf (*n* = 8).

Bleeding sap rate per plant can be separated into root number and function per unit root number (Morita & Abe, 2002). It is well established that in monocots, root emergence is associated with a regular pattern per phytomer (Nemoto et al., 1995), with approximately four roots per leaf in sugarcane (Miyazato, 1965). Therefore, we used leaf number as an indicator of root number. Under severe drought conditions (plots 3 and 4), bleeding sap rate per plant was partially determined based on the root number, but this was not the case under well-watered conditions (Figure 2e). Across all treatments, a strong relationship was detected between total water uptake and bleeding sap rate per leaf (an indicator of root function per unit root number), explaining 94.6% to 99.8% of the variation (Figure 2f).

At the same time, the mean leaf area ratio (LAR) explained 13.7% of the variation in mean RGR (Figure 2b), as also confirmed by specific leaf area (SLA) (Figure S3). Notably, no significant correlation was observed between NAR and LAR (*R^2^* < 0.001, Figure 2c). Therefore, improving NAR through enhanced root water uptake could be an effective strategy for increasing sugarcane productivity under drought conditions without negatively impacting biomass allocation to the leaves.

### Experiment 2: Enhanced Root Growth and Photosynthesis Promoted Drought Tolerance During Drought and Recovery in Harunoogi

In the pot experiment, the interaction between root function and photosynthesis was further investigated using the drought-sensitive cultivar NiTn18 and drought-tolerant Harunoogi. Plants were exposed to 12 days of drought stress followed by 13 days of rewatering (Figure S2b). Soil VWC in the pots ranged from 0.30 to 0.35 m^3^ m^−3^ under well-watered conditions and declined to 0.08 m^3^ m⁻^3^ during drought (Figure S2b). After drought treatment, leaf relative water content decreased from 98.54 ± 0.17% to 50.68 ± 2.17% in NiTn18 and from 98.49 ± 0.25% to 51.56 ± 6.38% in Harunoogi, with no significant difference between the two cultivars. Under well-watered conditions, shoot growth and root growth were comparable between the two cultivars (Figure 3a–c). However, after the drought and recovery phases, Harunoogi, exhibited a significantly greater leaf dry weight, stem dry weight, and leaf area (Figure 3a, 4a). With regard to root traits, sugarcane is characterized by two root types: sett roots, which emerge first from the seed setts and support early growth; and shoot roots, which are nodal roots that emerge from the base of the new shoot and develop into the main, deeper root system for water and nutrient uptake (Smith et al., 2005). In this experiment, after rewatering, Harunoogi displayed a significantly greater shoot root dry weight and shoot root number than NiTn18 (Figure 3b, c, right), whereas no significant differences were observed in sett root traits between the two cultivars (Figure 3b, c, left).

**Figure 3.**
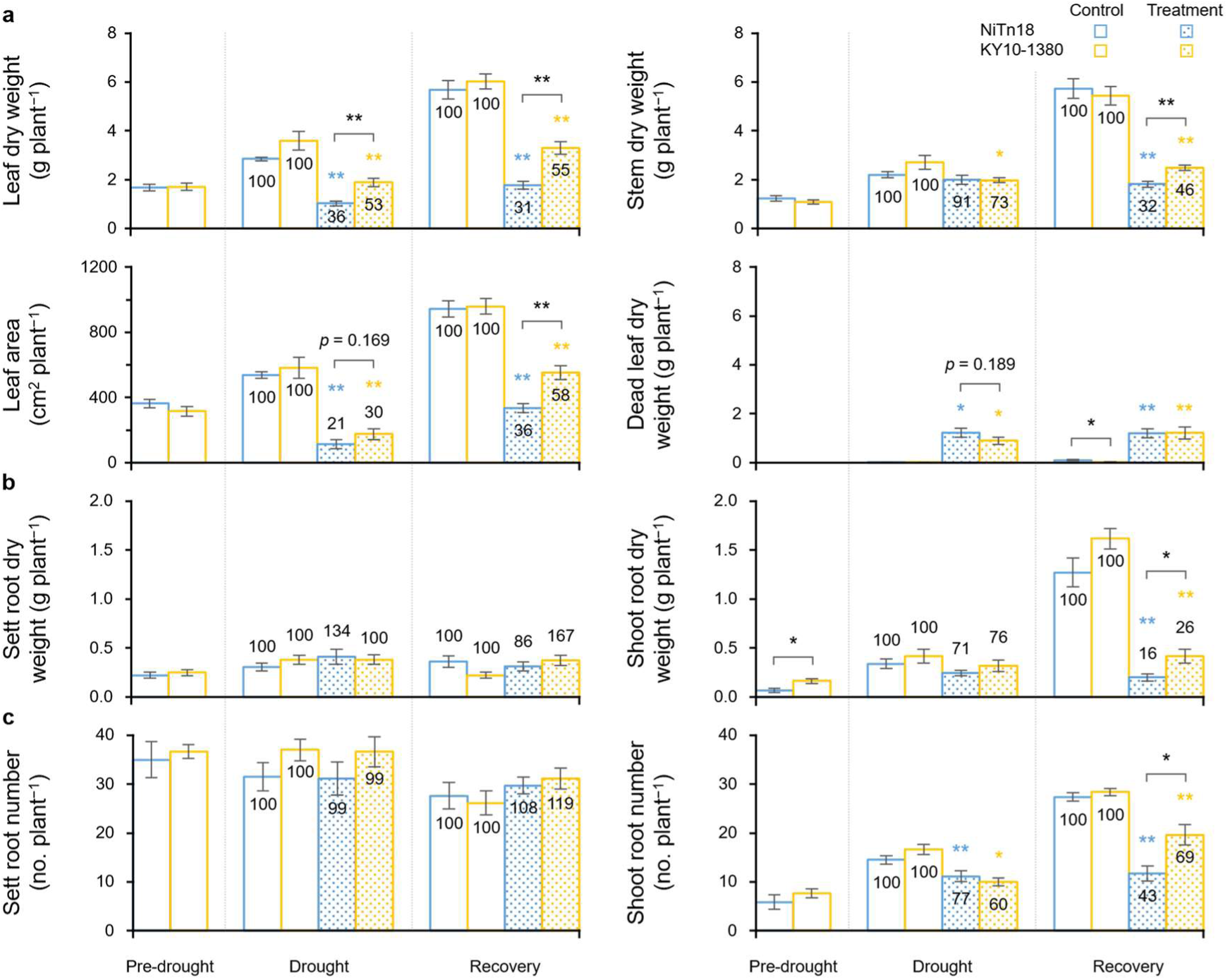
Shoot and root growth in cultivars NiTn18 and Harunoogi. **a** Dry weights of leaves, stem, and dead leaves, and leaf area; **b** root dry weight; and **c** root number. Error bars represent SEM (*n* = 7–8). Significant differences between cultivars (black) or treatments (colored) are denoted as \*\**p* < 0.01 or *0.05 based on Student’s *t*-test. The values in each column represent relative growth (%) under treatment compared to the control.

**Figure 4.**
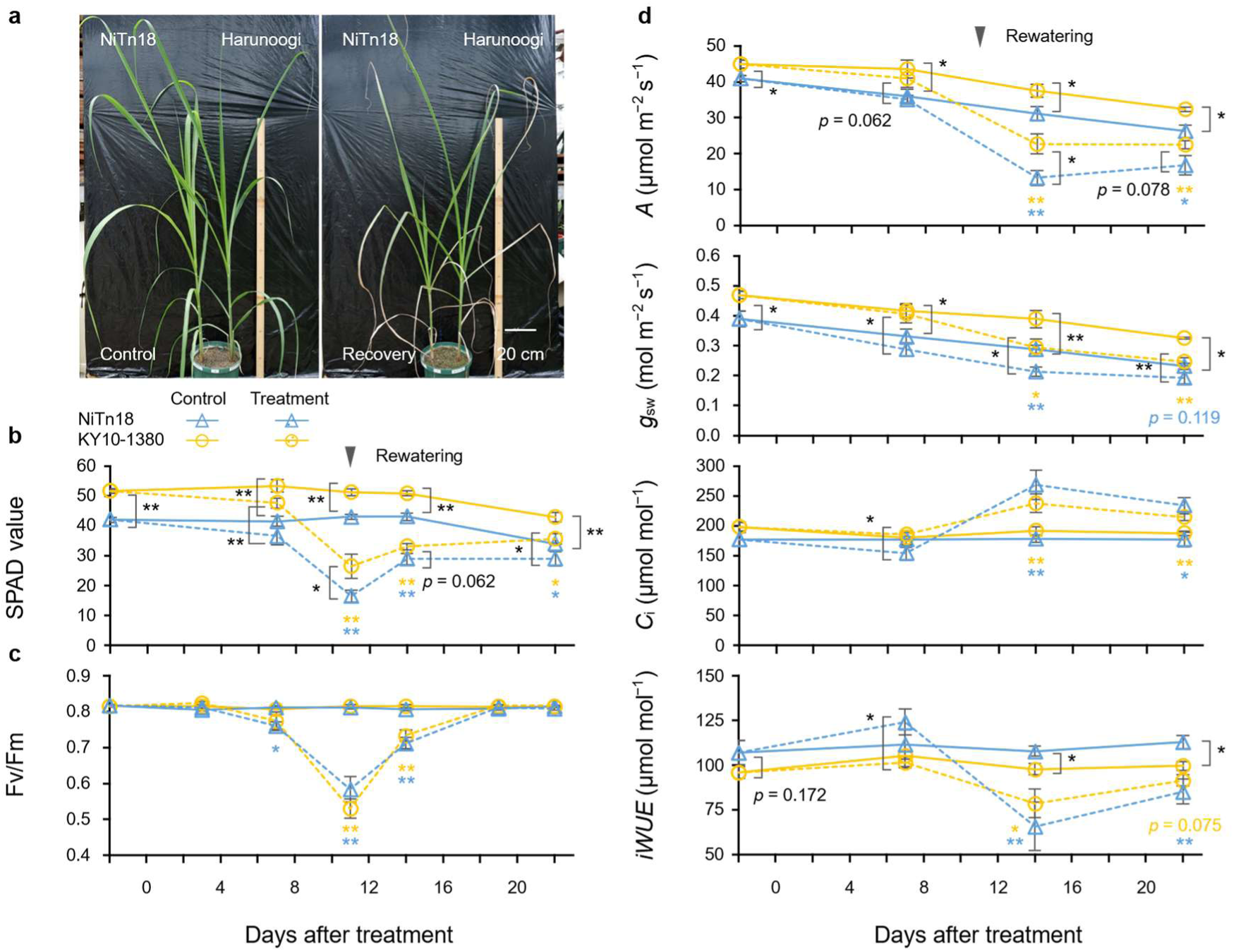
Leaf gas exchange in cultivar NiTn18 and Harunoogi during drought and recovery. **a** Growth performance after rewatering; **b** leaf chlorophyll content (SPAD value) in mature leaves; **c** maximum quantum yield of PSII (Fv/Fm) in the same leaves; and **d** steady-state net photosynthetic assimilation of CO_2_ (*A*), stomatal conductance (*g*_sw_), internal CO_2_ concentration (*C*_i_), and intrinsic water-use efficiency (*iWUE*) under a PPFD of 2000 µmol m^−2^ s^−1^. Error bars represent SEM (*n* = 5–8). Significant differences between cultivars (black) or treatments (colored) are denoted as \*\**p* < 0.01 or *0.05 based on Student’s *t*-test.

Continuous leaf measurements revealed that SPAD values (chlorophyll content) declined in both cultivars as the drought progressed but recovered after rewatering (Figure 4b). At the same time, compared with NiTn18, Harunoogi consistently maintained significantly higher chlorophyll *a* and *b* and total carotenoid contents under both control and treatment conditions (Figure S4a, b). The maximum quantum yield of PSII (Fv/Fm) showed a similar trend with no significant differences between cultivars, indicating that their maximum PSII activities were comparable (Figure 4c). Furthermore, Harunoogi exhibited higher stomatal conductance (*g*_sw_) and net CO_2_ assimilation rate (*A*) under both control and drought conditions as well as during recovery (Figure 4d).

Collectively, these findings indicate that Harunoogi’s superior drought tolerance is associated with enhanced shoot root growth, sustained leaf expansion, and improved carbon fixation via photosynthesis.

### Experiment 3: PSI Activity and Stomatal Conductance Played a Crucial Role in Maintaining Photosynthesis Under Drought Stress

To elucidate the photochemical mechanisms underlying the response to drought, we simultaneously measured the chlorophyll fluorescence of PSI and PSII during drought and recovery (Figure S2c). Fv/Fm followed a pattern similar to that observed in Experiment 2, with values for both cultivars declining during drought and recovering upon rewatering (Figure 5a). Notably, Harunoogi exhibited a significantly higher maximum amplitude of the reaction center chlorophyll of PSI (P700) (Pm) before and throughout the treatment (Figure 5b). Drought stress caused a greater decline in Pm than in Fv/Fm, with Pm requiring twice as long to recover after rewatering (Figure 5a, b). As a decrease in Pm indicates photoinhibition of PSI (Kono et al., 2017), sustained PSI activity is likely a key factor contributing to the superior *A* observed in Harunoogi under drought conditions (Figure 4d).

**Figure 5.**
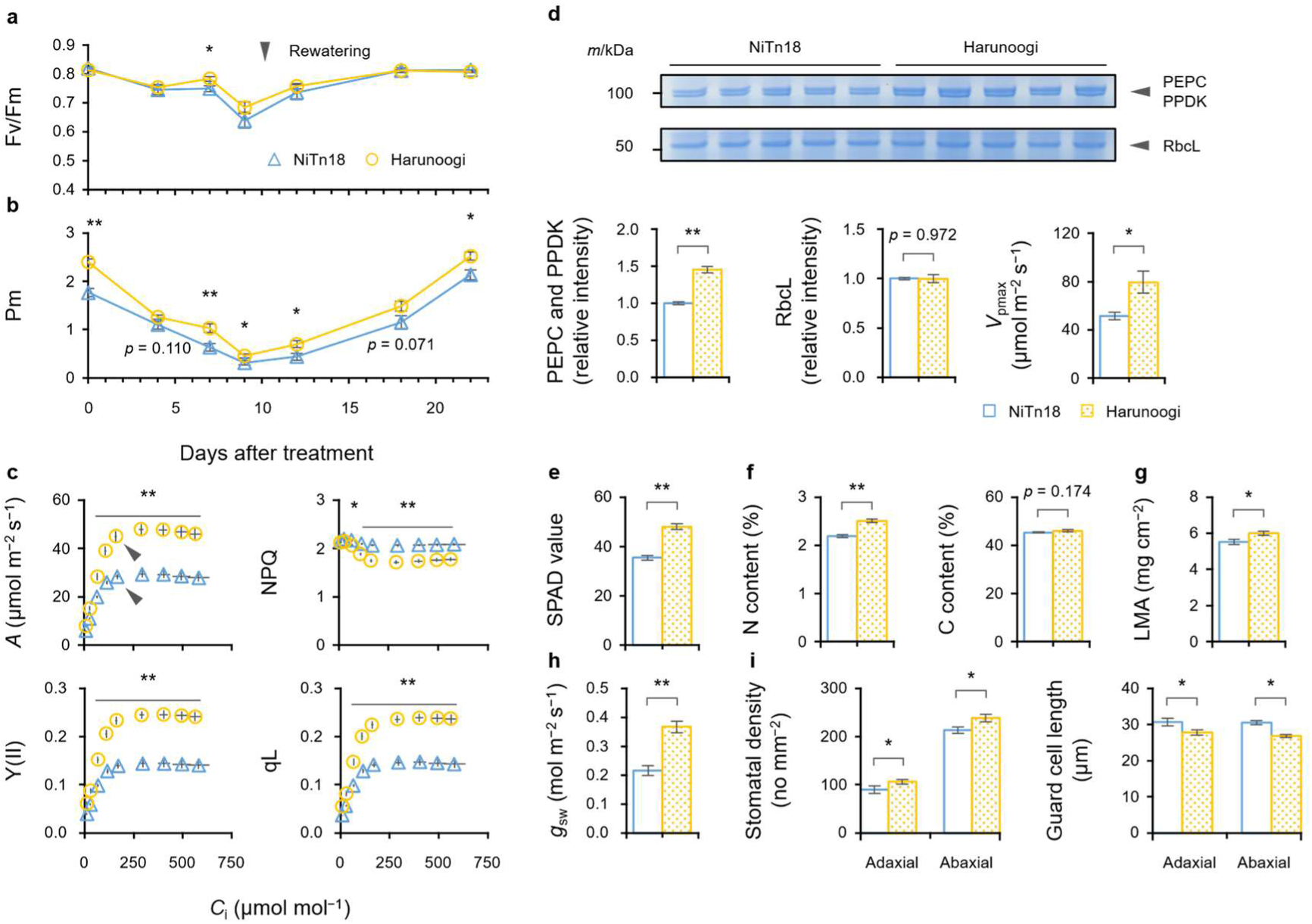
Photosynthetic capacity of cultivars NiTn18 and Harunoogi, and photoinhibition of PSI and PSII during drought and recovery. **a** Maximum quantum yield of PSII (Fv/Fm) in mature leaves. **b** Maximum amplitude of P700 (Pm) in the same leaves (*n* = 8). **c** CO_2_ response of net photosynthetic assimilation of CO_2_ (*A*), quantum yield of PSII (Y(II)), non-photochemical quenching (NPQ), and fraction of open PSII reaction centers (qL) under a PPFD at 2000 µmol m^−2^ s^−1^ before treatment (*n* = 8). Arrows indicate *A* under ambient CO_2_ concentration. **d** SDS-PAGE analysis of phospho*enol*pyruvate carboxylase (PEPC), pyruvate, phosphate dikinase (PPDK), and the large subunit of Rubisco (RbcL) before treatment. Relative signal intensities were quantified using the Gel Analyzer tool in ImageJ/Fiji software (*n* = 5). The apparent maximum rate of carboxylation by PEPC (*V*_pmax_) was calculated from *A*/*C*_i_ curves. **e** SPAD value (*n* = 8); **f** N and C contents; **g** leaf mass per area (LMA) before treatment (*n* = 10); **h** steady-state stomatal conductance (*g*_sw_) under a PPFD at 2000 µmol m^−2^ s^−1^ (*n* = 8); and **i** guard cell length and stomatal density in mature leaves before treatment (*n* = 6). Error bars represent SEM. Significant differences between cultivars are denoted as \*\**p* < 0.01 or *0.05 based on Student’s *t*-test.

The *A*/*C*_i_ curve revealed that *A* under ambient [CO_2_] fell within the transitional range between enzyme limitation and electron transport limitation (von Caemmerer, 2021) in well-watered sugarcane plants (Figure 5c). Although *A*/*C*_i_ curves were not measured under drought, a previous study conducted under similar conditions suggests that the transition point is unaffected by drought (Dinh et al., 2017). Under unsaturated *C*_i_, Harunoogi exhibited a superior electron sink capacity downstream of PSII, as indicated by the higher qL observed (Figure 5c). This was likely supported by the significantly higher *g*_sw_ associated with higher stomatal density (Figure 5h, i) and the higher apparent maximum rate of carboxylation by phospho*enol*pyruvate carboxylase (PEPC) (*V*_pmax_) linked to elevated PEPC, and pyruvate, phosphate dikinase (PPDK) contents (Figure 5d). Above the transitional *C*_i_, higher SPAD values (Figure 5e) were probably associated with improved light harvesting and electron transport in Harunoogi (Y(II), Figure 5c). Increases in C_4_ enzyme activity and chlorophyll content per leaf area were linked to enhanced nitrogen assimilation and increased leaf thickness (Figure 5f, g). Overall, these findings suggest that Harunoogi’s drought tolerance is driven by coordinated regulation of *g*_sw_, enzymatic activity, and photochemical efficiency, which collectively supported *A* under water-limited conditions.

## Discussion

### Robust Root Growth Supports Shoot Hydration and Drought Tolerance in Harunoogi

Under drought conditions, RGR was primarily determined by NAR, depending on both root number and efficiency (Figure 2a, d–f). Harunoogi developed significantly more shoot roots and exhibited a higher shoot root dry weight than NiTn18 after drought and rewatering (Figure 3b, c, right), whereas sett root traits were similar between cultivars (Figure 3b, c, left). In addition, shoot root dry weight was significantly lower in NiTn18 than in Harunoogi before drought, in line with previous findings showing that NiTn18 has a lower proportion of shoot root biomass relative to total root biomass compared to other commercial cultivars (Fukuzawa et al., 2008). This higher shoot root number and root biomass probably contributed to Harunoogi’s enhanced root water absorption under drought and recovery (Figure 1d). The higher leaf nitrogen content in Harunoogi (Figure 5f) also suggests that more nitrogen was acquired through root water uptake (Sakaigaichi et al., 2007), which is consistent with the function of shoot roots under drought (Smith et al., 2005).

Harunoogi may have inherited its robust root traits from its wild ancestor *Saccharum spontaneum* (Hattori et al., 2019), which shares deep-rooting and drought-tolerant traits with other wild relatives such as *Erianthus arundinaceus* (Lakshmanan & Robinson, 2013; Augustine et al., 2015; Terajima et al., 2023). For instance, an interspecific hybrid between *S. officinalum* and *S. spontaneum* was shown to recover its root length and biomass post-drought more rapidly than commercial sugarcane cultivars (Chanaphai et al., 2023), which is in line with our findings. These results support the value of wild germplasm for broadening the sugarcane genetic base and improving root function under drought.

### Sustained Photosynthesis Enhances Biomass Production under Drought

Across drought treatments, RGR was more correlated with NAR than with LAR (Figure 2a, b), which is consistent with previous field studies (Singh & Rao, 1987; Ramesh, 2000). Harunoogi maintained a higher net CO_2_ assimilation rate (*A*) than NiTn18 during drought and recovery (Figure 4d), which probably contributed to its higher NAR. This result also aligns with previous studies showing that maintaining photosynthesis under drought is important for root and shoot biomass production in sugarcane and other plants (Dinh et al., 2017; Vadez et al., 2024; Katsuhama et al., 2025).

Although *A* under well-watered conditions was higher in Harunoogi than in NiTn18, it did not lead to a significantly greater shoot dry mass (Figure 3a). Given that these two cultivars exhibited similar root traits under well-watered conditions (Figure 3b, c and 1d), it is possible that it was Harunoogi’s lower intrinsic water-use efficiency (*iWUE*) due to high *g*_sw_ (Figure 4d) that caused dehydration without additional growth. A similar situation has been reported in *Arabidopsis thaliana* and rice mutants with constitutively open stomata (Kusumi et al., 2017; Kimura et al., 2020). Furthermore, NiTn18 has steeper leaf angles than Harunoogi (Hattori et al., 2019; Terajima et al., 2010), which may enhance light absorption at the canopy level and contribute to whole-plant photosynthesis.

In summary, water and nitrogen uptake by the root system supports leaf expansion and photosynthesis, while carbon fixed in the leaves supports root growth (N. G. Inman-Bamber et al., 2012; Vadez et al., 2024). Owing to both its superior shoot root growth and enhanced water uptake, Harunoogi exhibited drought tolerance through sustained leaf photosynthesis.

### PSI Stability Contributes to Sustained Photosynthesis Under Drought

The maintenance of PSI activity per leaf area, indicated by the maximum amplitude of the reaction center chlorophyll of PSI (P700) (Pm) (Klughammer & Schreiber, 2008), underscores Harunoogi’s capacity to sustain *A* under drought stress (Figure 4d, 5b). In general, suppression of carbon fixation induced by abiotic stress causes over-reduction of PSI (Yamori & Shikanai, 2016). Electrons from the over-reduced PSI can in turn reduce O_2_, which generates reactive oxygen species (ROS) that damage the iron-sulfur centers of PSI, leading to its photoinhibition (Kono et al., 2024; Yamori & Shikanai, 2016). In rice, severe hyperosmotic stress was shown to lead to the over-reduction of P700, resulting in decreases in Pm, Fv/Fm, *A,* and biomass (Wada et al., 2019). Another study revealed that tobacco plants overexpressing the sugarcane *ScPetC* gene exhibited less chlorophyll degradation induced by oxidative damage than wild type plants, which was associated with higher Pm and Fv/Fm under drought (Silva et al., 2024). In Harunoogi, the higher chlorophyll *a* content per leaf area (or SPAD value) and fraction of open PSII reaction centers (qL) were associated with enhanced Pm, enabling higher PSII quantum yield (Y(II)) and *A* (Figure 5b, c and Figure S4).

Because PSI photoinhibition can lead to the over-reduction of plastoquinone, it may also over-reduce PSII, resulting in its photoinhibition (Kono et al., 2024). In the present study, Harunoogi, compared to NiTn18, maintained significantly higher Fv/Fm and Pm levels at 7 days after drought stress (Figure 5a, b), which suggests that enhanced PSI activity contributed to PSII protection. Notably, Pm declined earlier and recovered more slowly than Fv/Fm during treatment, indicating that PSI is more susceptible to drought under natural light conditions (Figure 5a, b). This is possibly due to the turnover of PSI being much slower than that of PSII once the system gets damaged and the fact that PSI recovery is further delayed under natural fluctuating light (Kono et al., 2024).

Overall, prolonged PSI photoinhibition may be a limiting factor for photosynthesis during recovery, further inhibiting biomass production in NiTn18 after rewatering (Figure 3a, 4d).

### Possible Biochemical Mechanisms Underlying PSI Stability under Drought

Given the vulnerability of PSI to oxidative damage when electron acceptors are limited, its sustained activity in Harunoogi suggests enhanced photoprotection and/or electron sink capacity mostly via carbon fixation (Yamori & Shikanai, 2016; Kono et al., 2024).

Harunoogi exhibited significantly higher *V*_pmax_ before drought (Figure 5d), which confirms a higher electron sink capacity through carbon fixation (Figure 5c). The high *g*_sw_ under drought (Figure 5h) may have helped maintain CO_2_ supply as a substrate to sustain enzymatic activity during the stressed period, and further prevented ROS generation by providing a sufficient electron sink to the photosystems (Yamori, 2016). Previous studies have shown that drought affects PEPC, PPDK, and related enzymes in sugarcane (Du et al., 1996; Sales et al., 2021). Prolonged stress can irreversibly damage biochemical functions and lead to a reduction in *iWUE* (Sales et al., 2021). In this study, Harunoogi showed slightly higher *iWUE* under drought than NiTn18 did due to a higher *A* relative to *g*_sw_ (Figure 4d), which suggests sustained carbon fixation. In addition, the fact that Harunoogi had higher total carotenoid contents (Figure S4b), which help dissipate excess energy through NPQ and ROS scavenging (Yamori & Shikanai, 2016; Goto et al., 2022), is indicative of greater photoprotection. Similar protective responses, including maintained chlorophyll content associated with higher ROS scavenging activity, have been observed in other drought-tolerant sugarcane genotypes (Endres et al., 2019; Silva et al., 2024).

Overall, sustained *g*_sw_, enzymatic activity, and photosynthetic pigment contents in Harunoogi likely contributed to maintaining Pm and eventually *A* during drought. Further studies on the balance among light harvesting, electron transport, and CO_2_ fixation under stress conditions—including both photosystems, C₃ and C₄ enzyme activities, and antioxidant systems—may help clarify the molecular basis of Harunoogi’s drought tolerance.

### Harnessing Wild Relatives for Breeding Climate-Resilient Sugarcane Cultivars

Harunoogi’s superior root and leaf functions highlight the potential of interspecific hybridization to expand the genetic background of commercial sugarcane and improve abiotic stress tolerance. Wild relatives such as *E. arundinaceus* and *S. spontaneum* offer valuable traits, including deep rooting and adaptability to marginal environments (Lakshmanan & Robinson, 2013; Terajima et al., 2023). Wild relatives have been considered valuable resources for improving photosynthesis in other crops. For example, in tomatoes, wild relatives adapted to diverse natural habitats showed higher daily integrated photosynthesis due to faster stomatal responses under fluctuating light conditions (Yoshiyama et al., 2024).

With the expected increase in atmospheric [CO_2_] in the future, breeding cultivars with a lower *g*_sw_ without reducing *A* could improve the water use efficiency of C_3_ and C_4_ crops (Leakey et al., 2019; Tanigawa et al., 2024). In sorghum, a 45% reduction in stomatal density achieved via synthetic EPF expression increased *iWUE* without compromising *A*, enhancing drought avoidance (Ferguson et al., 2024). Similarly, *E. arundinaceus* which has a lower *g*_sw_ than sugarcane due to reduced stomatal density, exhibits a higher *iWUE* under moderate drought conditions (Takaragawa & Wakayama, 2024). Furthermore, some *S. spontaneum* accessions demonstrate improved *iWUE* than commercial cultivar NCo310, due to higher *A* relative to *g*_sw_, which is associated with greater chlorophyll content and NADP-malic enzyme activity (Nose et al., 1994). Combined with our findings, the above results support the use of wild germplasm to improve root function and photosynthetic performance in sugarcane under abiotic stress.

## Conclusion

This study identified up to ninefold genetic variation in root water uptake among eight sugarcane cultivars under severe drought, showing a strong positive correlation between bleeding sap rate and net assimilation rate. By thoroughly comparing drought-tolerant cultivar Harunoogi and drought-sensitive NiTn18, we showed that enhanced shoot root growth is critical for maintaining water and nutrient supply under drought and recovery. Harunoogi also exhibited a consistently higher CO_2_ assimilation rate, which was supported by high stomatal conductance, chlorophyll content, and maximum amplitude of the reaction center chlorophyll of PSI. This is one of the first reports focusing on the interactions between the photosynthetic electron transport system (PSI and PSII) and leaf gas exchange in sugarcane under drought stress. Improving both root function and leaf photosynthesis may increase sugarcane yield potential and drought tolerance, assisting in addressing the challenges posed by future climate change.

## Acknowledgments

We thank Dr. Ryuta Terada, Kotaro Makino (Kagoshima University), Dr. Masaru Kono, and Dr. Ichiro Terashima (The University of Tokyo) for their technical support and insightful discussions regarding chlorophyll fluorescence measurements.

## Author Contributions

N.K., S.Y., and J.S. designed the study. N.K. and T.S. carried out the experiments and analyzed the data. N.K. wrote the initial manuscript draft. All authors reviewed the results and participated in revising the manuscript.

## Funding Details

This work was supported by Japan Science and Technology Agency (JST) under JST SPRING (JPMJSP2108 to N.K.).

## Disclosure statement

The authors report there are no competing interests to declare.

## Data Availability

Supporting data are available from the corresponding author on request.

---

**Supplementary Figure S1.**
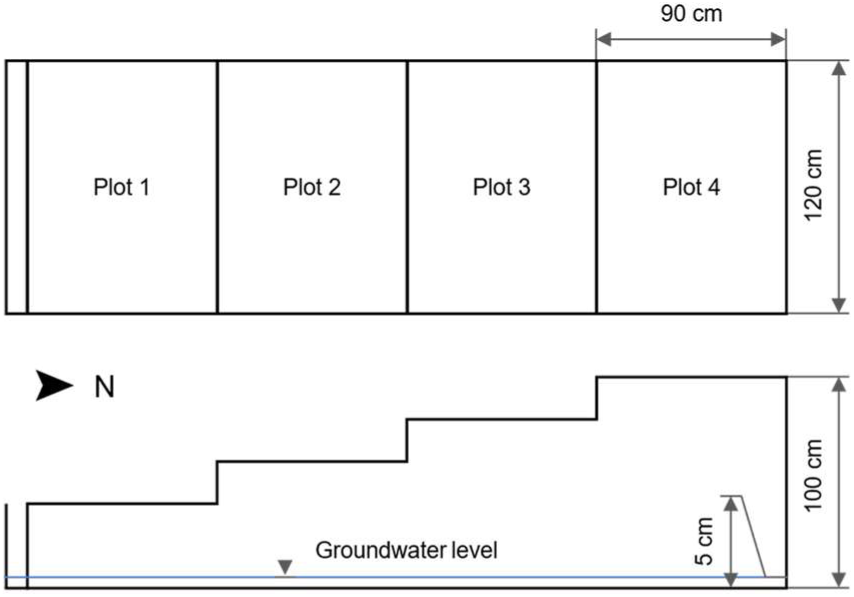
Diagram illustrating the sloped field used for Experiment 1. Soil volumetric water content was measured at 10 cm and 30 cm below the soil surface.

**Supplementary Figure S2.**
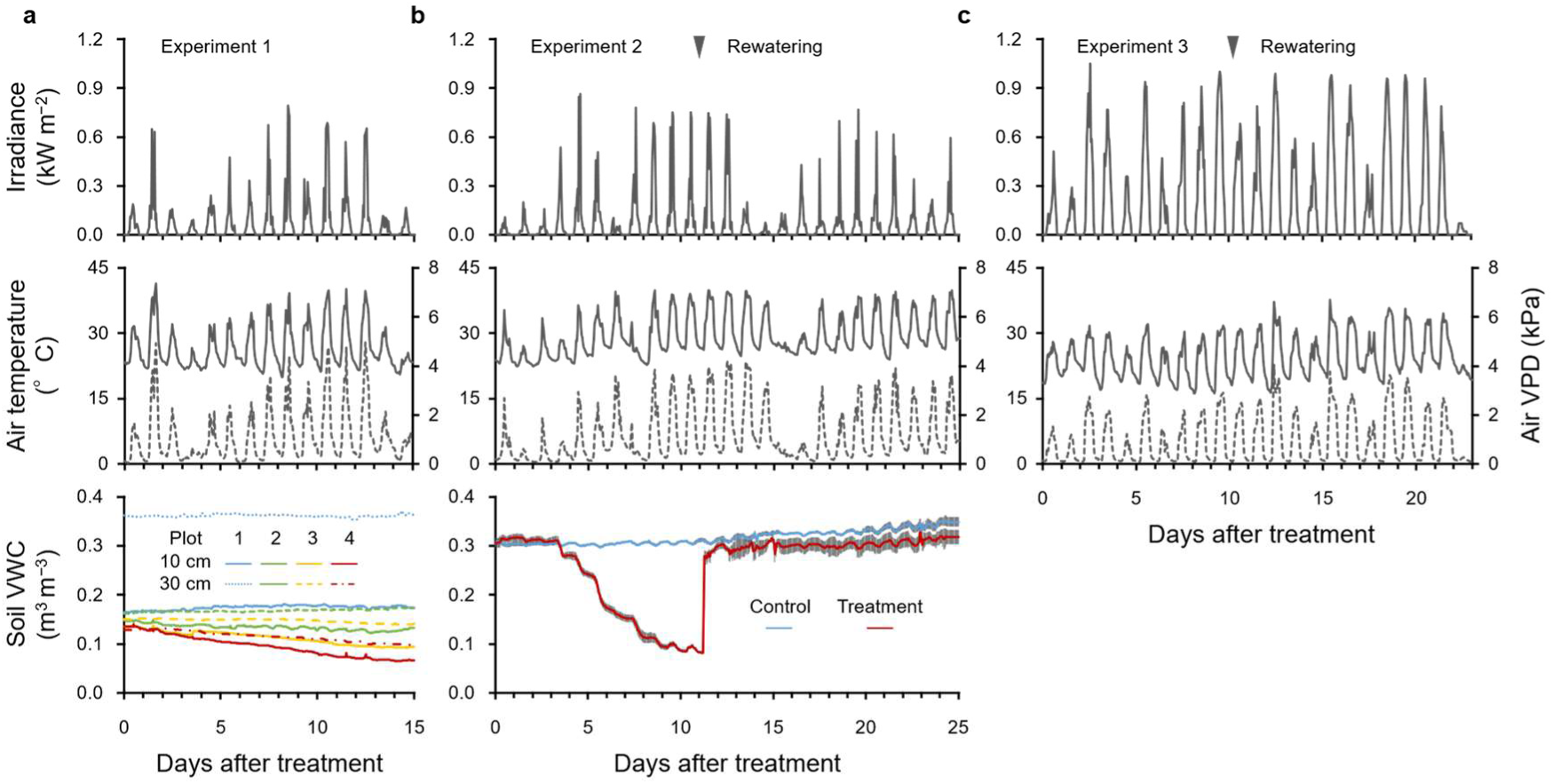
Solar radiation, atmospheric conditions, and soil water status during Experiments 1 (a), 2 (b), and 3 (c). Air vapor pressure deficit (VPD) was calculated from air temperature and relative humidity. Soil volumetric water content (VWC) was measured in Experiment 1 (*n* = 2) and Experiment 2 (*n* = 3). Error bars represent SEM. Arrows indicate the dates when watering was resumed.

**Supplementary Figure S3.**
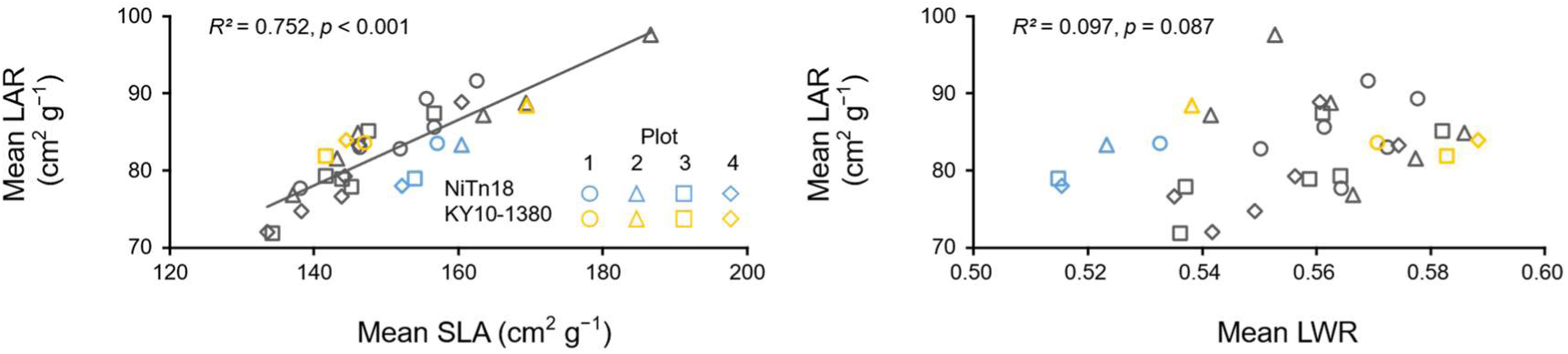
Relationship between mean leaf area ratio (LAR) and both mean specific leaf area (SLA) and mean leaf weight ratio (LWR) under four water conditions. Indices were calculated using mean shoot dry mass and leaf area from 5–8 biological replicates before and after treatment (*n* = 32).

**Supplementary Figure S4.**
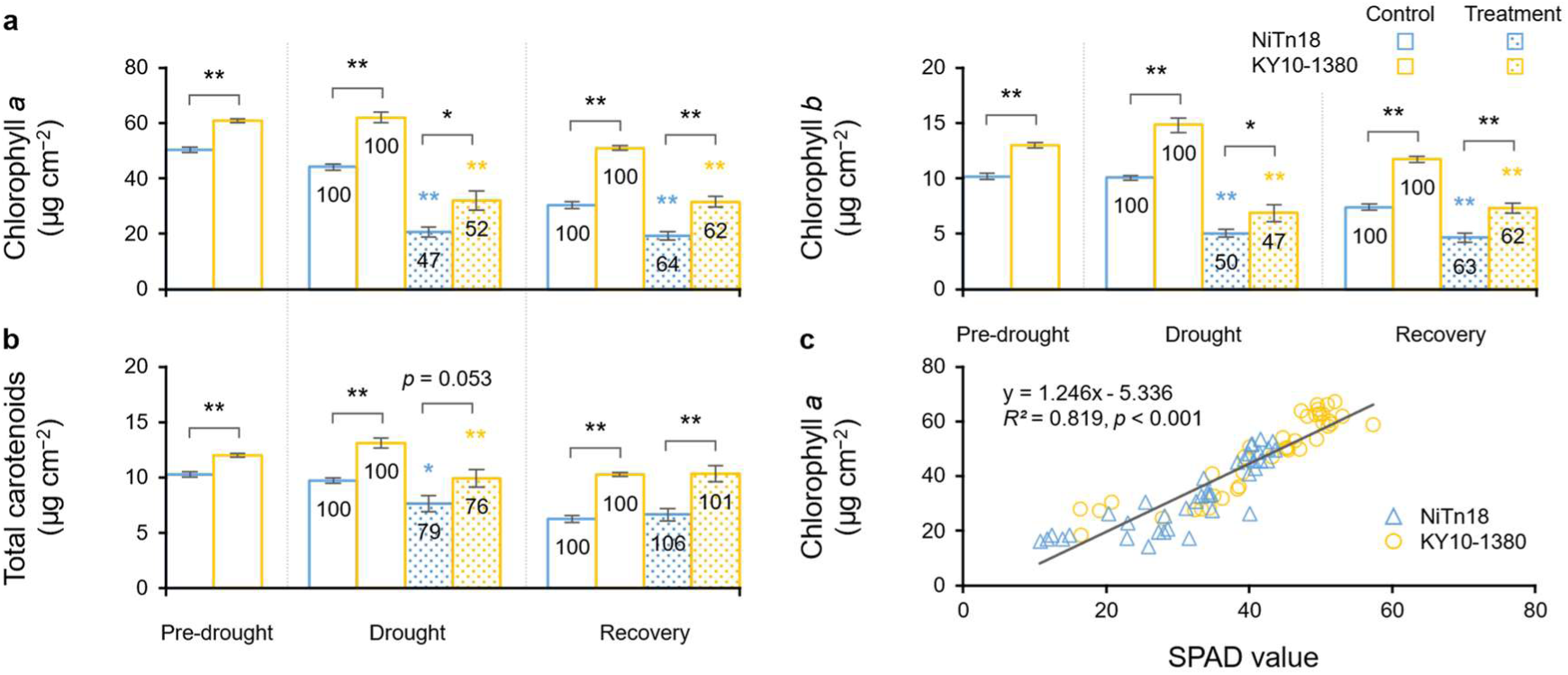
Chlorophyll and carotenoid contents in mature leaves. **a** Chlorophyll *a* and *b*; **b** total carotenoids. Error bars represent SEM (*n* = 7–8). Significant differences between cultivars (black) or treatments (colored) are denoted as \*\**p* < 0.01 or *0.05 based on Student’s *t*-test. The values in each column represent relative contents (%) measured in treated plants compared to the control. **c** Relationship between SPAD value and chlorophyll *a* content throughout the Experiment 2 (*n* = 78).

**Supplementary Figure S5.**
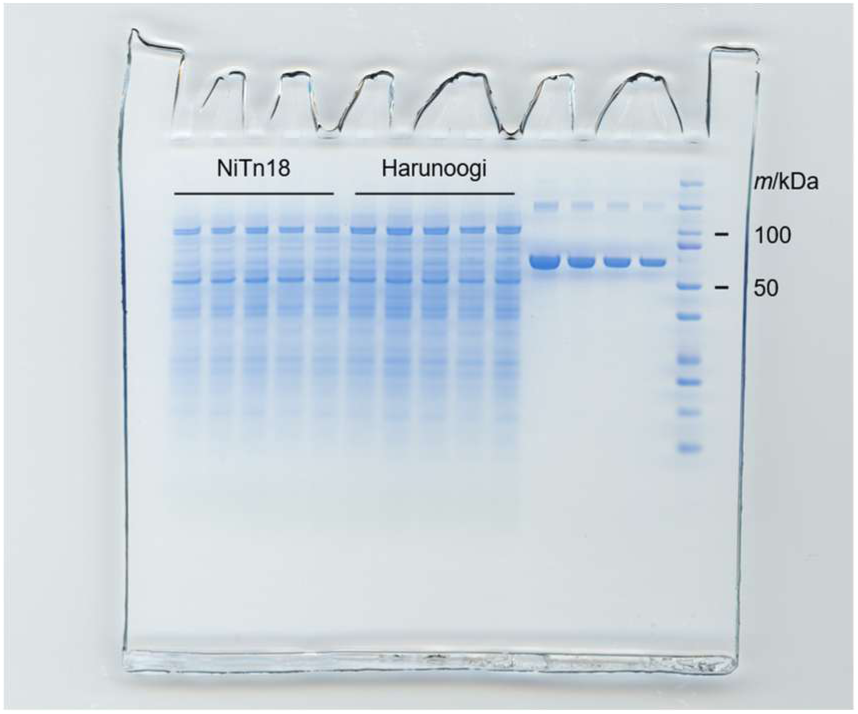
Uncropped gel image obtained from SDS-PAGE analysis, whose results are shown in Figure 5.

## Notes

### Competing Interest Statement

The authors have declared no competing interest.

## References

Augustine, S. M., Syamaladevi, D. P., Premachandran, M. N., Ravichandran, V., & Subramonian, N. (2015). Physiological and molecular insights to drought responsiveness in *Erianthus* spp. Sugar Tech, 17(2), 121–129. 10.1007/s12355-014-0312-7

Basnayake, J., Jackson, P. A., Inman-Bamber, N. G., & Lakshmanan, P. (2015). Sugarcane for water-limited environments. Variation in stomatal conductance and its genetic correlation with crop productivity. Journal of Experimental Botany, 66(13), 3945–3958. 10.1093/jxb/erv194

Chanaphai, P., Jongrungklang, N., Puangbut, D., & Songsri, P. (2023). Response of photosynthetic and root traits of sugarcane genotypes under drought and recovery conditions. Sugar Tech, 25(5), 1102–1114. 10.1007/s12355-023-01288-7

Cursi, D. E., Hoffmann, H. P., Barbosa, G. V. S., Bressiani, J. A., Gazaffi, R., Chapola, R. G., Junior, A. R. F., Balsalobre, T. W. A., Diniz, C. A., Santos, J. M., & Carneiro, M. S. (2022). History and current status of sugarcane breeding, germplasm development and molecular genetics in Brazil. Sugar Tech, 24(1), 112–133. 10.1007/S12355-021-00951-1/TABLES/6

de Almeida Silva, M., Jifon, J. L., Sharma, V., da Silva, J. A. G., Caputo, M. M., Damaj, M. B., Guimarães, E. R., & Ferro, M. I. T. (2011). Use of physiological parameters in screening drought tolerance in sugarcane genotypes. Sugar Tech, 13(3), 191–197. 10.1007/s12355-011-0087-z

Dinh, T. H., Watanabe, K., Takaragawa, H., Nakabaru, M., & Kawamitsu, Y. (2017). Photosynthetic response and nitrogen use efficiency of sugarcane under drought stress conditions with different nitrogen application levels. Plant Production Science, 20(4), 412–422. 10.1080/1343943X.2017.1371570

Du, Y. C., Kawamitsu, Y., Nose, A., Hiyane, S., Murayama, S., Wasano, K., & Uchida, Y. (1996). Effects of water stress on carbon exchange rate and activities of photosynthetic enzymes in leaves of sugarcane (*Saccharum* Sp.). Functional Plant Biology, 23(6), 719–726. 10.1071/pp9960719

Endres, L., dos Santos, C. M., Silva, J. V., Barbosa, G. V. de S., Silva, A. L. J., Froehlich, A., & Teixeira, M. M. (2019). Inter-relationship between photosynthetic efficiency, Δ13C, antioxidant activity and sugarcane yield under drought stress in field conditions. Journal of Agronomy and Crop Science, 205(5), 433–446. 10.1111/jac.12336

FAO. (2024). World food and agriculture – Statistical yearbook 2024. FAO. https://openknowledge.fao.org/handle/20.500.14283/cd2971en

Ferguson, J. N., Schmuker, P., Dmitrieva, A., Quach, T., Zhang, T., Ge, Z., Nersesian, N., Sato, S. J., Clemente, T. E., & Leakey, A. D. B. (2024). Reducing stomatal density by expression of a synthetic epidermal patterning factor increases leaf intrinsic water use efficiency and reduces plant water use in a C_4_ crop. Journal of Experimental Botany, 75(21), 6823–6836. 10.1093/jxb/erae289

Fukuzawa, Y., Kawamitsu, Y., Komiya, Y., & Ueno, M. (2008). Biomass production characteristics of sugarcane at initial growth stage. Japanese Journal of Crop Science, 77(1), 54–60. 10.1626/jcs.77.54

Goto, K., Yabuta, S., Tamaru, S., Ssenyonga, P., Emanuel, B., Katsuhama, N., & Sakagami, J.-I. (2022). Root hypoxia causes oxidative damage on photosynthetic apparatus and interacts with light stress to trigger abscission of lower position leaves in *Capsicum*. Scientia Horticulturae, 305, 111337. 10.1016/j.scienta.2022.111337

Hattori, T., Terajima, Y., Sakaigaichi, T., Terauchi, T., Tarumoto, Y., Adachi, K., Hayano, M., Tanaka, M., Ishikawa, S., Umeda, M., Matsuoka, M., & Sugimoto, A. (2019). High ratoon yield sugarcane cultivar “Harunoogi” developed for Kumage region by using an interspecific hybrid between a commercial cultivar and *Saccharum spontaneum* L. Journal of the NARO Research and Development, 2019(2), 21–44. 10.34503/naroj.2019.2_21

Hu, C., Elias, E., Nawrocki, W. J., & Croce, R. (2023). Drought affects both photosystems in *Arabidopsis thaliana*. New Phytologist, 240(2), 663–675. 10.1111/nph.19171

Hunt, R., Causton, D. R., Shipley, B., & Askew, A. P. (2002). A modern tool for classical plant growth analysis. Annals of Botany, 90(4), 485–488. 10.1093/AOB/MCF214

Inman-Bamber, G., Moore, P. H., & Botha, F. C. C. (2013). Sugarcane yields and yield-limiting processes. Sugarcane: Physiology, Biochemistry, and Functional Biology, 579–600. 10.1002/9781118771280.CH21

Inman-Bamber, N. G., Lakshmanan, P., & Park, S. (2012). Sugarcane for water-limited environments: Theoretical assessment of suitable traits. Field Crops Research, 134, 95–104. 10.1016/J.FCR.2012.05.004

IPCC. (2022). Climate change and land: IPCC special report on climate change, desertification, land degradation, sustainable land management, food security, and greenhouse gas fluxes in terrestrial ecosystems. 1–896. 10.1017/9781009157988

Jackson, P. A. (2005). Breeding for improved sugar content in sugarcane. Field Crops Research, 92(2–3), 277–290. 10.1016/J.FCR.2005.01.024

Jackson, P., Basnayake, J., Inman-Bamber, G., Lakshmanan, P., Natarajan, S., & Stokes, C. (2016). Genetic variation in transpiration efficiency and relationships between whole plant and leaf gas exchange measurements in *Saccharum* spp. And related germplasm. Journal of Experimental Botany, 67(3), 861–871. 10.1093/JXB/ERV505

Kato, Y., Abe, J., Kamoshita, A., & Yamagishi, J. (2007). Varietal Differences in Stem Diameter and Rooting Number of Phytomers in Conjunction with Root System Development of Field-Grown Rice (*Oryza sativa* L.). Plant Production Science, 10(3), 357–360. 10.1626/pps.10.357

Katsuhama, N., Sakoda, K., Kimura, H., Shimizu, Y., Sakai, Y., Nagata, K., Abe, M., Terashima, I., & Yamori, W. (2025). PROTON ATPASE TRANSLOCATION CONTROL 1-mediated H^+^-ATPase translocation boosts plant growth under drought by optimizing root and leaf functions. PNAS Nexus, 4(5), pgaf151. 10.1093/pnasnexus/pgaf151

Kimura, H., Hashimoto-Sugimoto, M., Iba, K., Terashima, I., & Yamori, W. (2020). Improved stomatal opening enhances photosynthetic rate and biomass production in fluctuating light. Journal of Experimental Botany, 71(7), 2339–2350. 10.1093/jxb/eraa090

Klughammer, C., & Schreiber, U. (2008). Saturation Pulse method for assessment of energy conversion in PS I. PAM Application Note, 1, 11–14.

Kono, M., Oguchi, R., & Terashima, I. (2024). Photoinhibition of PSI and PSII in nature and in the laboratory: Ecological approaches. In U. Lüttge, F. M. Cánovas, M.-C. Risueño, C. Leuschner, & H. Pretzsch (Eds.), Progress in Botany Vol. 84 (pp. 241–292). Springer Nature Switzerland. 10.1007/124_2022_67

Kono, M., Yamori, W., Suzuki, Y., & Terashima, I. (2017). Photoprotection of PSI by far-red light against the fluctuating light-induced photoinhibition in *Arabidopsis thaliana* and Field-Grown Plants. Plant and Cell Physiology, 58(1), 35–45. 10.1093/pcp/pcw215

Kramer, D. M., Johnson, G., Kiirats, O., & Edwards, G. E. (2004). New fluorescence parameters for the determination of QA redox state and excitation energy fluxes. Photosynthesis Research, 79(2), 209–218. 10.1023/B:PRES.0000015391.99477.0d

Kusumi, K., Hashimura, A., Yamamoto, Y., Negi, J., & Iba, K. (2017). Contribution of the S-type anion channel SLAC1 to stomatal control and its dependence on developmental stage in rice. Plant and Cell Physiology, 58(12), 2085–2094. 10.1093/pcp/pcx142

Lakshmanan, P., & Robinson, N. (2013). Stress physiology: Abiotic stresses. *Sugarcane: Physiology*, Biochemistry, and Functional Biology, 411–434. 10.1002/9781118771280.CH16

Leakey, A. D. B., Ferguson, J. N., Pignon, C. P., Wu, A., Jin, Z., Hammer, G. L., & Lobell, D. B. (2019). Water use efficiency as a constraint and target for improving the resilience and productivity of C_3_ and C_4_ crops. Annual Review of Plant Biology, 70, 781–808. 10.1146/annurev-arplant-042817-040305

Mall, A. K., Misra, V., Pathak, A. D., & Srivastava, S. (2022). Breeding for drought tolerance in sugarcane: Indian perspective. Sugar Tech, 24(6), 1625–1635. 10.1007/S12355-021-01094-Z/TABLES/2

Miyazato K. (1965). Studies on the growth habit in an early stage of sugarcane. The science bulletin of the Division of Agriculture, Home Economics & Engineering, University of the Ryukyus, 12, 1–86.

Morita, S., & Abe, J. (2002). Diurnal and phenological changes of bleeding rate in lowland rice plants. Japanese Journal of Crop Science, 71(3), 383–388. 10.1626/jcs.71.383

Nemoto, K., Morita, S., & Baba, T. (1995). Shoot and root development in rice related to the Phyllochron. Crop Science, 35(1), 24–29. 10.2135/CROPSCI1995.0011183X003500010005X

Nose, A., Uehara, M., Kawamitsu, Y., Kobamoto, N., & Nakama, M. (1994). Variations in leaf gas exchange traits of *Saccharum* including feral sugarcane, *Saccharum spontaneum* L. Japanese Journal of Crop Science, 63(3), 489–495. 10.1626/jcs.63.489

OECD/FAO. (2024). OECD-FAO agricultural outlook 2024-2033. 10.1787/4C5D2CFB-EN

Ramesh, P. (2000). Effect of different levels of drought during the formative phase on growth parameters and its relationship with dry matter accumulation in sugarcane. Journal of Agronomy and Crop Science, 185(2), 83–89. 10.1046/j.1439-037x.2000.00404.x

Robertson, M. J., Inman-Bamber, N. G., Muchow, R. C., & Wood, A. W. (1999). Physiology and productivity of sugarcane with early and mid-season water deficit. Field Crops Research, 64(3), 211–227. 10.1016/S0378-4290(99)00042-8

Sakaigaichi, T., Morita, S., Abe, J., & Yamaguchi, T. (2007). Diurnal and Phenological Changes in the Rate of Nitrogen Transportation Monitored by Bleeding in Field-GrownRice Plants (*Oryza sativa* L.). Plant Production Science, 10(3), 270–276. 10.1626/PPS.10.270

Sales, C. R. G., Wang, Y., Evers, J. B., & Kromdijk, J. (2021). Improving C_4_ photosynthesis to increase productivity under optimal and suboptimal conditions. Journal of Experimental Botany, 72(17), 5942–5960. 10.1093/jxb/erab327

Schindelin, J., Arganda-Carreras, I., Frise, E., Kaynig, V., Longair, M., Pietzsch, T., Preibisch, S., Rueden, C., Saalfeld, S., Schmid, B., Tinevez, J. Y., White, D. J., Hartenstein, V., Eliceiri, K., Tomancak, P., & Cardona, A. (2012). Fiji: An open-source platform for biological-image analysis. Nature Methods, 9(7), 676–682. 10.1038/nmeth.2019

Silva, C. R. L., de Souza, C. B., dos Santos, C. M., Floreste, B. S., Zani, N. C., Hoshino-Bezerra, A. A., Bueno, G. C., Chagas, E. B. R., & Menossi, M. (2024). The sugarcane *ScPetC* gene improves water-deficit and oxidative stress tolerance in transgenic tobacco plants. Agronomy, 14(7), 7. 10.3390/agronomy14071371

Singh, S., & Rao, P. N. G. (1987). Varietal differences in growth characteristics in sugar cane. The Journal of Agricultural Science, 108(1), 245–247. 10.1017/S0021859600064327

Smith, D. M., Inman-Bamber, N. G., & Thorburn, P. J. (2005). Growth and function of the sugarcane root system. Field Crops Research, 92(2–3), 169–183. 10.1016/j.fcr.2005.01.017

Sun, H., Shi, Q., Liu, N.-Y., Zhang, S.-B., & Huang, W. (2023). Drought stress delays photosynthetic induction and accelerates photoinhibition under short-term fluctuating light in tomato. Plant Physiology and Biochemistry, 196, 152–161. 10.1016/j.plaphy.2023.01.044

Takaragawa, H., & Wakayama, M. (2024). Responses of leaf gas exchange and metabolites to drought stress in different organs of sugarcane and its closely related species *Erianthus arundinaceus*. Planta, 260(4), 90. 10.1007/s00425-024-04508-w

Tanigawa, K., Yuchen, Q., Katsuhama, N., Sakoda, K., Wakabayashi, Y., Tanaka, Y., Sage, R., Lawson, T., & Yamori, W. (2024). C_4_ monocots and C_4_ dicots exhibit rapid photosynthetic induction response in contrast to C_3_ plants. Physiologia Plantarum, 176(4), e14431. 10.1111/ppl.14431

Terajima, Y., Hattori, T., Shimatani, M., Sato, M., Takaragawa, H., Sakaigaichi, T., Umeda, M., Naito, T., & Irei, S. (2022). Sugarcane Breeding and Supporting Genetics Research in Japan. Sugar Tech, 24(1), 134–150. 10.1007/S12355-020-00930-Y/FIGURES/8

Terajima, Y., Sugimoto, A., Matsuoka, M., Ujihara, K., Sakaigaichi, T., Fukuhara, S., Maeda, H., Katsuta, Y., Oka, M., Shimoda, S., Mizumoto, F., Higashi, T., Shikura, F., Urabe, K., Hayashi, T., Sato, M., Yoshida, N., Fukui, K., Hidaka, N., & Ueno, K. (2010). New sugarcane cultivar “NiTn18” with excellent ratooning ability in mulch-free cultivation. Bulletin of the National Agricultural Research Center for Kyushu Okinawa Region, 54, 23–41.

Terajima, Y., Sugimoto, A., Tippayawat, A., Irei, S., & Hayashi, H. (2023). Root distribution and fibre composition of intergeneric F1 hybrid between sugarcane and *E. arundinaceus*. Field Crops Research, 297, 108920. 10.1016/j.fcr.2023.108920

Uga, Y., Sugimoto, K., Ogawa, S., Rane, J., Ishitani, M., Hara, N., Kitomi, Y., Inukai, Y., Ono, K., Kanno, N., Inoue, H., Takehisa, H., Motoyama, R., Nagamura, Y., Wu, J., Matsumoto, T., Takai, T., Okuno, K., & Yano, M. (2013). Control of root system architecture by *DEEPER ROOTING 1* increases rice yield under drought conditions. Nature Genetics, 45(9), 1097–1102. 10.1038/ng.2725

UNESCO. (2020). United Nations world water development report 2020: Water and climate change. UNESCO.

Vadez, V., Grondin, A., Chenu, K., Henry, A., Laplaze, L., Millet, E. J., & Carminati, A. (2024). Crop traits and production under drought. Nature Reviews Earth & Environment, 5(3), 211–225. 10.1038/s43017-023-00514-w

von Caemmerer, S. (2021). Updating the steady-state model of C_4_ photosynthesis. Journal of Experimental Botany, 72(17), 6003–6017. 10.1093/jxb/erab266

Wada, S., Takagi, D., Miyake, C., Makino, A., & Suzuki, Y. (2019). Responses of the photosynthetic electron transport reactions stimulate the oxidation of the reaction center chlorophyll of photosystem I, P700, under drought and high temperatures in rice. International Journal of Molecular Sciences, 20(9), Article 9. 10.3390/ijms20092068

Wang, X., Wang, H., Liu, S., Ferjani, A., Li, J., Yan, J., Yang, X., & Qin, F. (2016). Genetic variation in *ZmVPP1* contributes to drought tolerance in maize seedlings. Nature Genetics, 48(10), 1233–1241. 10.1038/ng.3636

Wang, X.-Q., Zeng, Z.-L., Li, Y.-Y., & Huang, W. (2024). Photoinhibition and photosynthetic regulation in fluctuating light under compound stresses of drought and heat. Physiologia Plantarum, 176(3), e14406. 10.1111/ppl.14406

Wellburn, A. R. (1994). The Spectral Determination of chlorophylls *a* and *b*, as well as total carotenoids, using various solvents with spectrophotometers of different resolution. Journal of Plant Physiology, 144(3), 307–313. 10.1016/S0176-1617(11)81192-2

Yamori, W. (2016). Photosynthetic response to fluctuating environments and photoprotective strategies under abiotic stress. Journal of Plant Research, 129(3), 379–395. 10.1007/s10265-016-0816-1

Yamori, W., Nagai, T., & Makino, A. (2011). The rate-limiting step for CO_2_ assimilation at different temperatures is influenced by the leaf nitrogen content in several C_3_ crop species. Plant, Cell & Environment, 34(5), 764–777. 10.1111/J.1365-3040.2011.02280.X

Yamori, W., & Shikanai, T. (2016). Physiological functions of cyclic electron transport around photosystem I in sustaining photosynthesis and plant growth. Annual Review of Plant Biology, 67, 81–106. 10.1146/annurev-arplant-043015-112002

Ye, H., Roorkiwal, M., Valliyodan, B., Zhou, L., Chen, P., Varshney, R. K., & Nguyen, H. T. (2018). Genetic diversity of root system architecture in response to drought stress in grain legumes. Journal of Experimental Botany, 69(13), 3267–3277. 10.1093/jxb/ery082

Yoshiyama, Y., Wakabayashi, Y., Mercer, K. L., Kawabata, S., Kobayashi, T., Tabuchi, T., & Yamori, W. (2024). Natural genetic variation in dynamic photosynthesis is correlated with stomatal anatomical traits in diverse tomato species across geographical habitats. Journal of Experimental Botany, 75(21), 6762–6777. 10.1093/jxb/erae082

